# Bacteriophage Sf14 survival under simulated gastrointestinal conditions

**DOI:** 10.64898/2026.09.24.754092

**Authors:** Giselle Hernandez, Hailey R. Kerns, Sarah M. Doore

## Abstract

*Shigella flexneri* is an enteric pathogen responsible for causing shigellosis, a type of bacillary dysentery of significant global health concern. The increasing prevalence of antibiotic resistance has driven the search for alternative treatments, including bacteriophage (phage) therapy. However, the effectiveness of orally administered phages for enteric pathogens depends on their ability to survive passage through the gastrointestinal tract. In this study, the ability of bacteriophage Sf14 to maintain infectivity was evaluated under simulated gastrointestinal conditions using a modified INFOGEST 2.0 *in vitro* digestion model. This model includes salivary, gastric, and intestinal fluids with physiologically relevant compositions of electrolytes, bile salts, and enzymes. Phages were incubated in each simulated phase at standard physiological temperature, then tested for infectivity using quantitative plaque assays. Results show that Sf14 remained stable during the short duration of the oral phase and for the duration of the intestinal phase. By contrast, survival in simulated gastric fluid required a pH of 2.75 or higher, with rapid and substantial loss of infectivity at pH 2.5 and below. These findings demonstrate that while Sf14 is stable in oral and intestinal environments, the gastric phase poses a critical barrier to phage survival. With *in vivo* fasting gastric pH ranging from 1.5 to 3, encapsulation would enable the most flexible oral phage therapy against enteric pathogens, but ingestion after eating or alongside antacids may be sufficient.

**Importance:** The bacterium *Shigella flexneri* causes shigellosis, a highly transmissible and potentially life-threatening intestinal infection in humans. It is becoming increasingly difficult to treat as antibiotic resistance rises. Lytic bacteriophages, viruses that specifically infect and destroy bacterial cells, offer a potential alternative to antibiotics. However, developing effective phage therapy requires not only identifying a phage that targets *S. flexneri*, but ensuring that sufficient viable phage particles survive the changing conditions of the gastrointestinal tract and reach the site of infection. This study identifies the acidic stomach environment as a major barrier to the oral delivery of bacteriophage Sf14 and demonstrates that small changes in gastric acidity can substantially alter its survival. Moreover, these findings emphasize the importance of considering physiological barriers during the preliminary stages of phage therapy development and may inform future dosing, formulation, and administration strategies for oral phage therapies for shigellosis.

## Introduction

*Shigella* are Gram-negative, facultatively intracellular bacteria that cause shigellosis, a type of bacillary dysentery (1). Globally, *Shigella* are estimated to be the second leading cause of diarrheal mortality, disproportionately affecting young children in regions with limited access to clean water, sanitation, health-care facilities, and health-care interventions. Among children under 5 years old, its annual toll has been estimated at 81,800 deaths and 7.34 million disability-adjusted life years (2). While the *Shigella* genus contains *S. boydii, S. dysenteriae, S. flexneri*, and *S. sonnei*, most of these cases can be attributed to the species *S. flexneri* (3). Following ingestion, *S. flexneri* targets the colonic epithelium, where it invades and grows within epithelial cells (4, 5). At the cellular level, this invasion is primarily mediated by a type III secretion system, which transfers effector proteins into host cells to promote bacterial uptake and survival. After entering the cell, *S. flexneri* escapes the phagocytic vacuole and spreads to adjacent cells, leading to inflammation, tissue damage, and the typical symptoms of dysentery as epithelial cells are damaged (4, 5). Antibiotics may be used to reduce the duration of bacterial excretion, but this treatment strategy has been hindered due to the rising resistance of *S. flexneri* to commonly used antibiotics (6). This has placed additional strain on the already limited healthcare systems of developing countries, reinforcing the need to develop alternative strategies, including vaccines or other antimicrobial approaches (7).

Lytic bacteriophages, which are viruses that infect and lyse specific bacterial hosts, have been proposed as a natural and effective intervention against a variety of bacterial pathogens (8). Bacteriophages are appealing alternatives since they can evolve alongside their host bacteria, their infection strategies are diverse, and they are abundant in nature. One type of bacteriophage that infects and lyses *S. flexneri* is phage Sf14, which represents a group of *S. flexneri-*infecting Moogleviruses (9). Sf14 can infect numerous serotypes of *S. flexneri* and could prove useful in the treatment or prevention of shigellosis. However, the effectiveness of bacteriophages in reducing bacterial populations can be influenced by environmental conditions, including temperature and pH, which may affect both bacterial activity and phage function (10, 11). Since oral delivery would be the most effective and accessible treatment for shigellosis, phages used for treatment must be able to survive the gastrointestinal tract. In this study, we evaluated whether bacteriophage Sf14 could remain infectious under simulated gastrointestinal conditions without additional protection, such as encapsulation. A modified version of the INFOGEST 2.0 protocol (12), adapted to exclude a food bolus, was used to generate simulated salivary, gastric, and intestinal fluids (**Figure 1)**. Phage infectivity was then assessed across these conditions via quantitative plaque assays. Results show that Sf14 retains full infectivity in salivary and intestinal fluids, and that most particles can also survive in the mid-to upper-level pH levels of the gastric phase. Since gastric pH rises to this range shortly after eating, this suggests that Sf14 may be a viable candidate for oral phage therapy with only modifications to delivery timing or methods.

**Figure 1.**
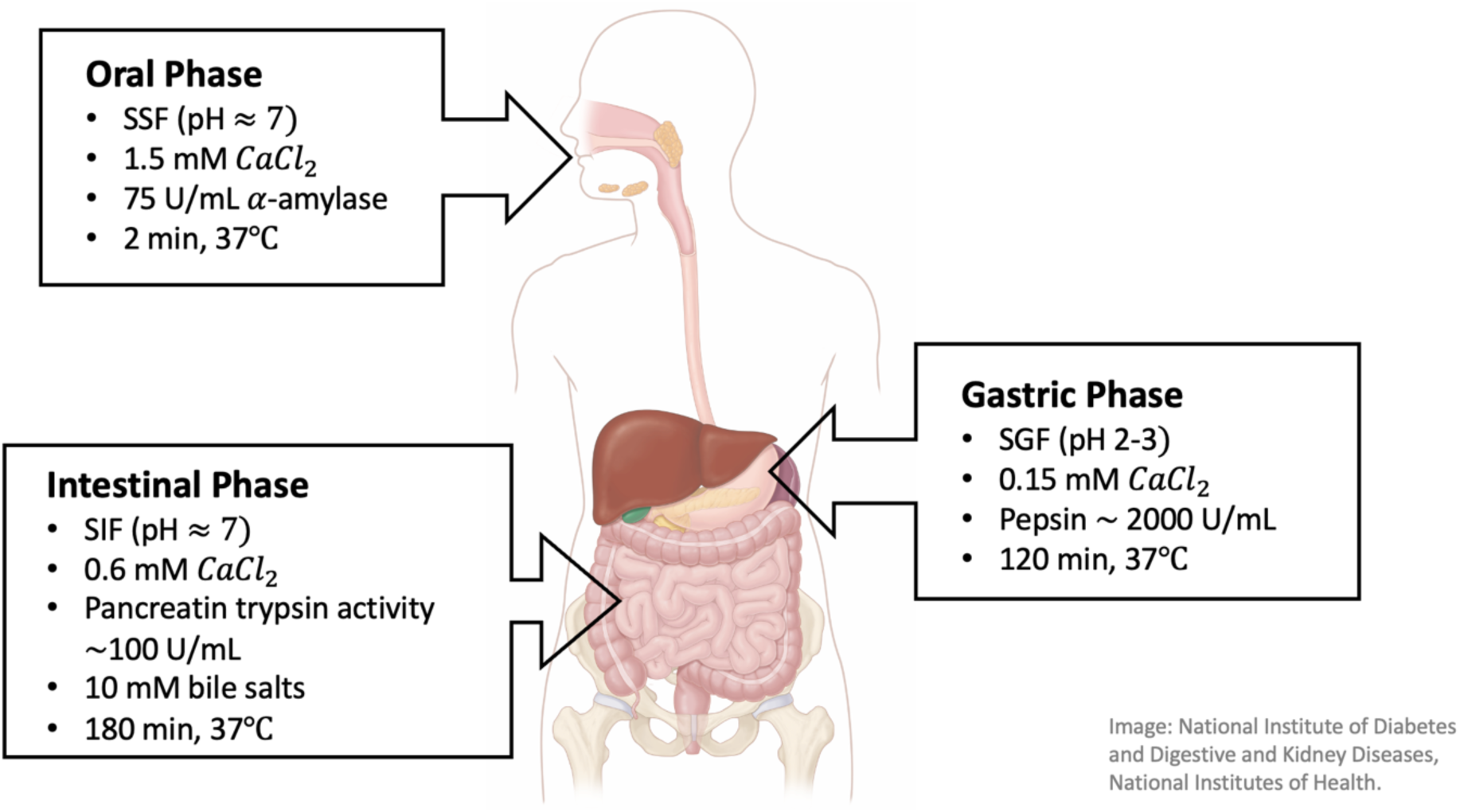
Target conditions for *in vitro* simulation of oral, gastric, and intestinal digestion of bacteriophage Sf14.

## Results and Discussion

### Oral Phase

During digestion, the oral phase is comparatively brief and mild. It consists of near-neutral pH, moderate ionic strength, and brief residence times: conditions under which many bacteriophages remain stable. Although a 2-minute oral phase is likely much longer than *in vivo* digestion, this was the duration tested based on standard recommendations (12, 13), with infectivity tested at 0, 0.5, 1, and 2 min. For Sf14, exposure to simulated salivary fluid (SSF) resulted in little to no detectable loss of infectivity across the full exposure period. Plaque counts obtained at 0, 0.5, 1, and 2 min were generally comparable, indicating the phage can survive the conditions tested here (**Figure 2**). This is consistent with the relatively mild physicochemical environment of the oral cavity, which is characterized by near-neutral pH, low proteolytic activity, and short residence times. Although α-amylase was included in the simulated salivary fluid, its enzymatic activity is specific to carbohydrate hydrolysis and is not expected to affect phage structural proteins or nucleic acids. While incubations of 5 min have been used for some static models (14), the lack of change observed here and overall short duration of the oral phase is not expected to pose a significant barrier to the phage.

**Figure 2.**
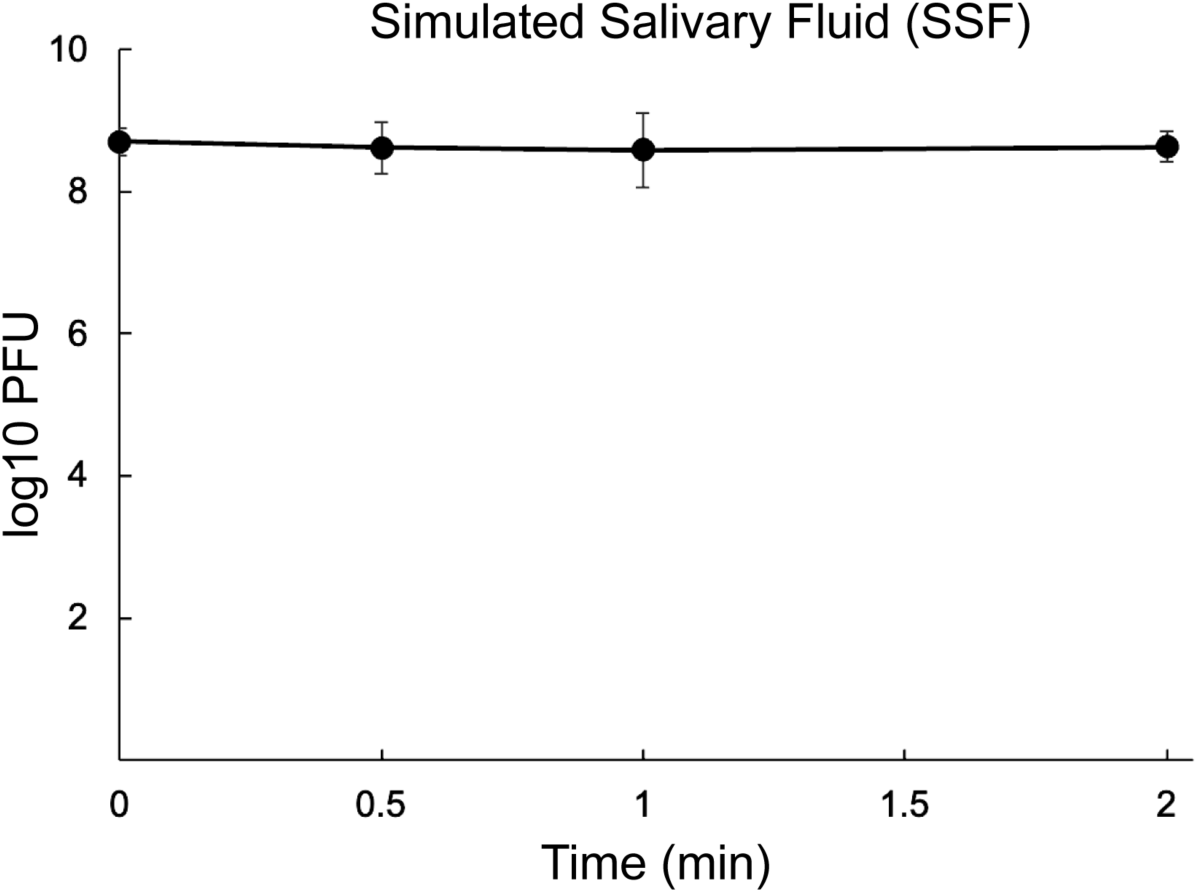
Infectivity of Sf14 over time when incubated in simulated salivary fluid (SSF) for up to 2 min.

### Acidic Conditions and the Gastric Phase

The stomach was anticipated to be the harshest digestive phase for Sf14 due to its strong acidity. *In vivo*, gastric pH varies frequently: it can decrease to values as low as pH 1.5 while fasting, then rise to 5.0 after food ingestion in adults (15) or up to 6.9 after liquid feeding in neonates (16). This suggests that static models using a constant pH of 3.0 may fail to accurately predict microbial survival (12). To determine an initial range of Sf14 survival under acidic conditions, trials were conducted across pH 2 – 3 in standard phage dilution buffer to account for physiological variability in stomach acidity.

Timepoints up to 120 min were included to distinguish rapid inactivation from progressive loss of infectivity over a physiologically relevant gastric exposure period. Gastric emptying occurs over several hours, with standardized clinical protocols assessing gastric retention up to 4 hr following meal ingestion (16). Although gastric residence varies with ingested contents, liquids generally empty more rapidly than solids, making a 120 min exposure period relevant to the liquid, food-free model used here. As shown in **Figure 3A**, only slight differences in highly acidic pH produced noticeably different infectivity outcomes. At pH 2.5 and below, Sf14 was inactivated within 5 min. At pH 2.75, plaques were variably detected at 2 hr but experienced an overall loss of at least 10^-5^. At pH 3.0, there was a loss of nearly 10^-2^ by 2 hr, representing a significantly slower rate of decay.

**Figure 3.**
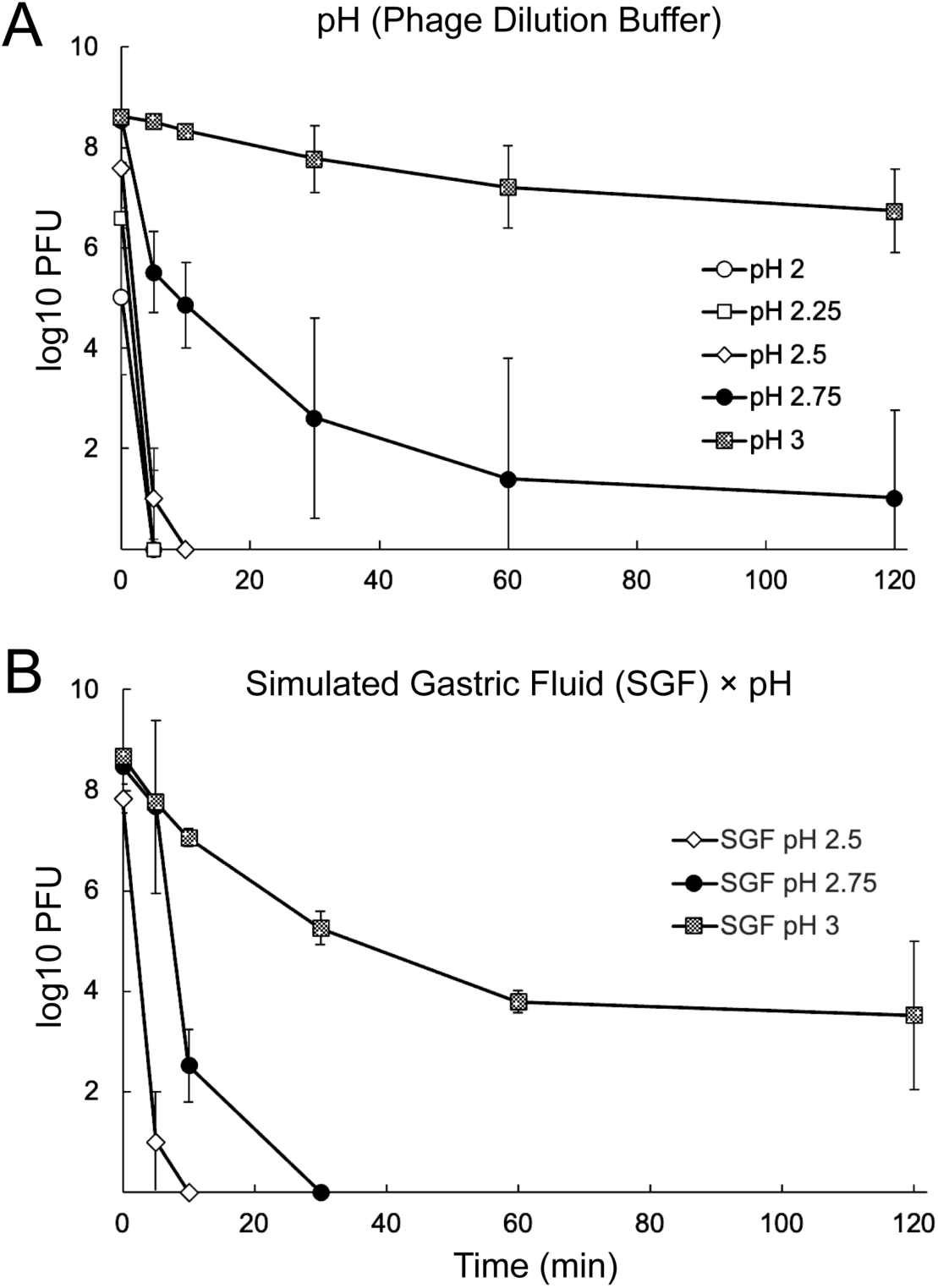
Infectivity of Sf14 over the course of 2 hrs when incubated at pH in the 2-3 range in **A**) standard phage dilution buffer or **B**) simulated gastric fluid (SGF).

Since the composition of gastric fluid is more complex than phage dilution buffer, we next tested simulated gastric fluid (SGF) at pH ranges from 2.5 to 3.0 (**Figure 3B**). As expected, Sf14 was inactivated within 5 min at SGF pH 2.5. In contrast to the experiments described above, virions were also inactivated at SGF pH 2.75 by 30 min. At pH 3.0, phages were still detectable at 2 hr but had experienced a loss of 10^-5^ to 10^-7^. Based on the decimal reduction time at pH 3.0, the maximum survival time of 10^8^ plaque forming units (PFU) would be approximately 3 – 3.5 hr. Conversely, if pH were the only variable, it would take approximately 3 hr at pH 2.75 to invariably reduce 10^8^ PFU below the detectable limit, but 11.5 hr at pH 3.0 to reach the same level of clearance.

These findings highlight the critical role of environmental conditions in determining phage viability during gastrointestinal transit. The acidic environment of the stomach, combined with digestive enzymes, is known to be a barrier for many phages. Intragastric pH is physiologically variable within and between individuals depending on numerous factors, including diet, medications, fasted gastric volume, sex, and circadian rhythm (17, 18). Parietal cells lining the stomach secrete hydrochloric acid with a pH of approximately 0.8, maintaining the highly acidic environment (19); however, food ingestion temporarily increases pH due to dilution and buffering of the food (20). One study involving young, healthy adults reported the median fasted gastric pH to be approximately 1.7; food administration temporarily increased participants’ gastric pH to a median peak of 6.7 before gradually returning to baseline within 2 hr (21). Additionally, food’s buffering effect may vary with meal composition, with high-volume fatty meals providing a buffering effect above pH 4 for 150 min (22).

Acidic conditions can destabilize viral capsid proteins and disrupt phage particle integrity, while the protease pepsin may further degrade exposed structural proteins (23). Together, these conditions likely contributed to the rapid inactivation of Sf14 particles observed in the experiment. Similar reductions in bacteriophage viability under acidic conditions have been reported in previous studies, which demonstrate that phages exhibit optimal stability and infectivity near neutral pH (approximately pH 7–8), while exposure to acidic conditions (pH < 6) results in substantial loss of viability and lytic activity due to structural damage and protein destabilization (24). However, the greater loss of Sf14 infectivity in SGF compared to pH-specific PDB suggests that acidity alone does not fully account for the rapid inactivation. When pH alone was considered, the time for 10^8^ PFU to degrade below detectable limits was significantly different when pH and SGF were combined. At pH 3.0 in standard phage buffer, Sf14 would have lasted nearly half a day; at pH 3.0 in SGF, it would only last 3 hr. The difference at pH 2.75 is even more stark, with survival of 3 hr at pH 2.75 in phage buffer compared to 15 min at pH 2.75 in SGF. While many phage characterization studies include pH in phage buffer, these results demonstrate that phage particle stability is a complex multifactorial interaction with the surrounding environment, with pH representing only one component.

*S. flexneri* is known to survive passage through the stomach by inducing acid resistance mechanisms under moderately acidic conditions, including structural modifications to the lipopolysaccharide that enhance its further resistance to extreme acidity (25, 26). Since phage Sf14 virions are not capable of similar response reactions, acidic stress likely destabilizes phage particles enough to expose structural proteins to proteases, leading to further degradation. Together, these factors contribute to irreversible damage to the phage particle and loss of infectivity. The gastric phase results also show that even small differences of 0.25 pH unit can substantially influence phage survival. *In vivo*, such variability may arise from factors such as food intake, buffering capacity, and individual physiological differences. Factors such as the timing and composition of food intake, using antacids and/or proton pump inhibitors, or a combination, may be sufficient to reduce stomach acidity enough to let virions pass through this phase without the need for additional encapsulation strategies. For oral administration of a Staphylococcal phage, treatment with proton pump inhibitors or eating yogurt facilitated passage into the small intestine, even when the phage particles were otherwise inactivated at by a 1 hr incubation at pH of 4 or below (27). Therefore, while the gastric environment represents a major barrier, it may not be uniformly restrictive under all conditions and could require only minor interventions.

### Intestinal Phase

Once past the stomach, the intestinal lumen provides a more neutral pH environment, ranging from 5 to 8 (15); however, there are numerous digestive components that could degrade phage particles, including enzymes and bile salts. The porcine pancreatin extract used here is a combination of digestive enzymes, including multiple proteases, lipases, nucleases, and amylase. In addition to enzymes, bile salts are amphipathic derivatives of cholesterol, having both negatively charged and hydrophobic regions that act as detergents to digest fats. Although Sf14 is not known to contain lipids, the negative charge of the bile salts could still degrade particles. We therefore wanted to determine whether the presence of pancreatin and bile salts in the simulated intestinal fluid would destabilize virions.

Bacteriophage Sf14 remained stable during exposure to simulated intestinal fluid over the full 3 hr incubation period (**Figure 4**). At the initial timepoint (t = 0), the mean phage titer was 4.9 ± 1.0 × 10⁸ PFU/mL (log₁₀ ≈ 8.7). A slight decrease in infectivity was observed by the end of the experiment, with mean titers of 4.6 ± 1.2 × 10⁸ PFU/mL at 1 hr and 3.4 ± 1.7 × 10⁸ PFU/mL at 3 hr (log₁₀ ≈ 8.7 and 8.5). These decreases were not statistically significant, indicating minimal loss of infectivity under intestinal conditions. Although the residence time in the small intestine ranges approximately 3 – 6 hr, Sf14 is likely resistant to the effects of bile salts and pancreatic enzymes at physiologically relevant concentrations. Although it does not produce its own digestive enzymes, the colon residence time varies from 16 hr to 29 hr (28), with some reports of over 50 hr (29). This duration is unlikely in the context of shigellosis and is unlikely to pose a barrier to particle stability: thus, the intestine appears to be a favorable environment for phage Sf14.

**Figure 4.**
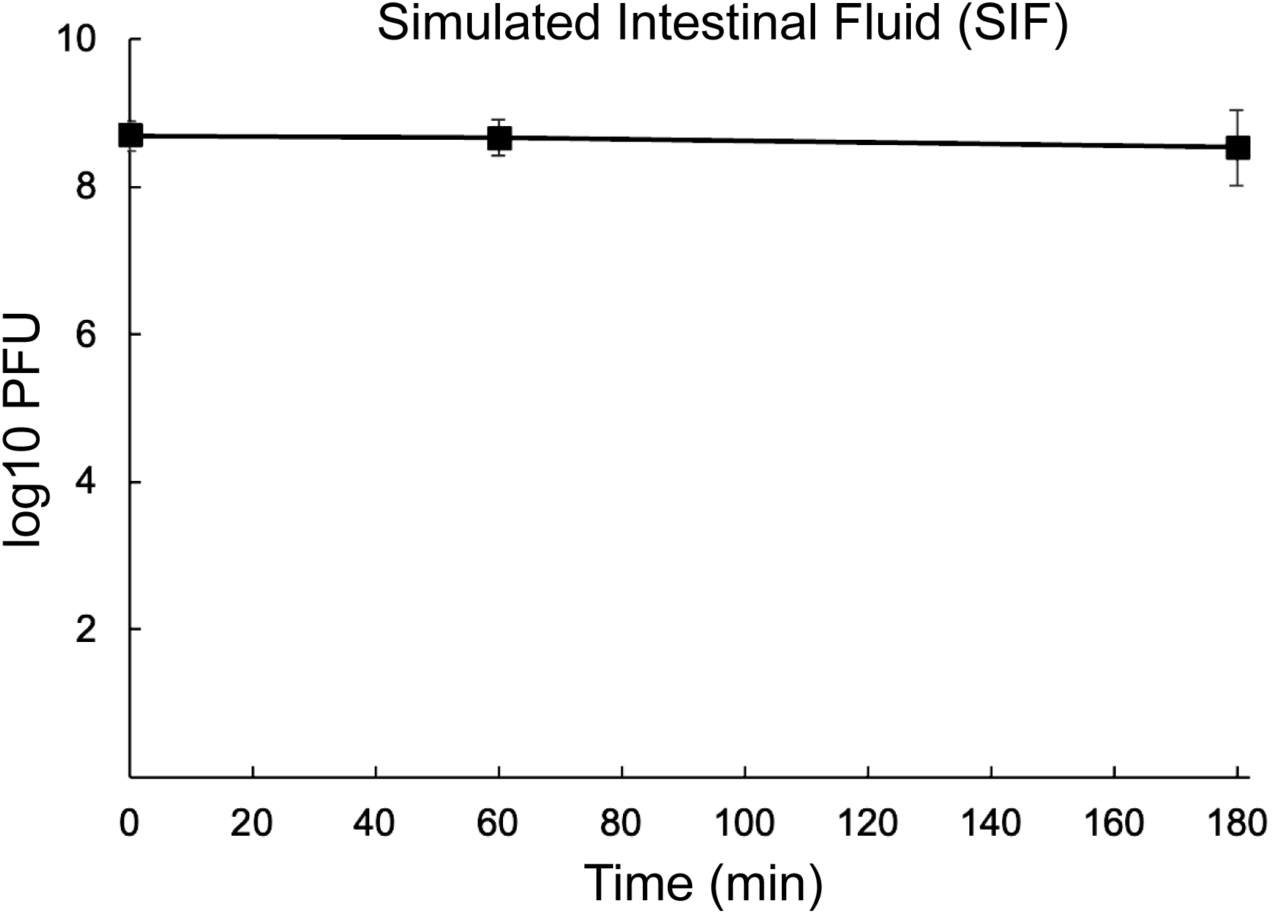
Infectivity of Sf14 over time when incubated in simulated intestinal fluid (SIF) for 3 hr.

### Conclusions

Using an *in vitro* simulated gastrointestinal model, our results demonstrate that bacteriophage Sf14 maintains infectivity under simulated human oral and intestinal conditions but is highly sensitive to the acidic conditions of the gastric environment. If at least some virions can transit the stomach, they are likely to remain viable in the region where *S. flexneri* colonization occurs. Bile salts play a regulatory role in *S. flexneri* physiology, promoting biofilm formation and bacterial aggregation in the small intestine, followed by dispersion and host cell interaction downstream (30). Depending on how soon after exposure the phage is ingested, this aggregation may either hinder phage access to individual cells or, conversely, facilitate localized phage amplification once infection is established within a cluster. Once *S. flexneri* reaches the colon and begins intracellular invasion and cell-to-cell spread, they become less accessible to the phage. While there is no evidence supporting intracellular killing by phages (31), phage treatment after the onset of symptoms has still been shown to reduce the duration and severity of disease (as reviewed in (32)).

Several limitations of this study should be considered. The use of a static *in vitro* digestion model does not fully replicate the dynamic nature of the gastrointestinal tract, where pH, enzyme concentrations, and transit times fluctuate continuously (12). Additionally, the absence of a food matrix may have led to an overestimation of phage inactivation, as food components can provide buffering effects that enhance microbial survival. The model also does not account for interactions with mucus layers, native gut microbiota, or the potential for bacterial aggregation and biofilm formation, which may influence phage stability and access to host bacteria. This work evaluates phage stability in the absence of host bacteria and therefore does not account for potential phage replication and amplification that may occur *in vivo* upon encountering *S. flexneri*. Furthermore, digestion phases were assessed independently rather than sequentially, which may not fully capture cumulative effects during gastrointestinal transit. However, other phages have been characterized using *in vitro* gastrointestinal conditions as part of initial characterization (34, 35). Others have examined complex *in vitro* systems as potential precursors or comparisons to *in vivo* studies (36, 37). Similar experiments using T4 have also identified how specific structural components affect particle stability in simulated gastrointestinal conditions (38).

Altogether, these results demonstrate that while Sf14 can persist in oral and intestinal environments, the gastric phase represents a critical limitation for therapeutic application. Future work may focus on developing protective delivery strategies, such as timing with food intake, encapsulation, or co-administration with buffering agents to improve phage survival during gastric transit. The use of an antacid just prior to phage addition has been sufficient for some phages (e.g. (34)). Increasing the initial delivery dose may also be useful in overcoming these harsher conditions. Incorporating more physiologically relevant models, including dynamic digestion systems or *in vivo* studies, will be essential for accurately predicting phage behavior and advancing the development of effective oral phage therapies targeting enteric pathogens such as *Shigella flexneri*.

## Materials & Methods

### Bacteria and Phage Methods

*S. flexneri* strain CFS100, an avirulent derivative of strain 2457T (33), was used as the bacterial host for bacteriophage Sf14 propagation and quantitative plaque assays (9). To produce high-titer phage stocks, LB broth was inoculated with a fresh overnight of CFS100 using a 2% inoculum. This was incubated at 37°C with shaking until the culture reached early log-phase growth, at which point approximately 10^7^ PFU were added to initiate infection. The infected culture was incubated for 3-4 hr, or until lysis. The lysate was centrifuged at 8,000 × g for 10 min to remove bacterial debris, then the supernatant was transferred to a new tube and centrifuged again at 13,500 × g for 45 min to pellet the phage. The phage pellet was gently resuspended in phage dilution buffer (PDB; 10 mM Tris HCl pH 7.6, 10 mM MgCl_2_) and stored at 4 °C. Stock titers were determined by plaque assay prior to the simulated digestion trials. The stock was diluted to create working phage suspension of approximately 10^9^ PFU per mL. Enumeration of viable particles was conducted using 0.1 mL of serial dilutions combined with 0.15 mL of overnight bacteria via soft agar overlay. All plates were incubated at 37°C overnight before quantification.

### Preparation of Simulated Digestive Electrolyte Solutions

Simulated salivary fluid (SSF), simulated gastric fluid (SGF), and simulated intestinal fluid (SIF) electrolyte solutions were prepared according to the INFOGEST 2.0 static *in vitro* digestion protocol (12). To ensure consistency across digestion phases, master salt stock solutions—KCl, KH₂PO₄, NaHCO₃, NaCl, MgCl₂·6H₂O, (NH₄)₂CO₃, and CaCl₂·2H₂O—were prepared at the defined molar concentrations listed in the INFOGEST 2.0 protocol (12; **Table 1**). These master stocks were used to prepare phase-specific electrolyte solutions, then stored at −20 °C.

**Table 1.** Components used in preparation of 1.25× Electrolyte Solutions (recipes are for 30 mL each). Final concentration refers to the final reaction mixture, not individual phase stocks.

| Stock Solutions | 1.25× SSF |  | 1.25× SGF |  | 1.25× SIF |  |
| --- | --- | --- | --- | --- | --- | --- |
|  | μL Stock | Final mM Concentration | μL Stock | Final mM Concentration | μL Stock | Final mM Concentration |
| **0.5 M KCl** | 1130 | 15.1 | 520 | 6.9 | 510 | 6.8 |
| **0.5 M KH<sub>2</sub>PO<sub>4</sub>** | 280 | 3.7 | 70 | 0.9 | 60 | 0.8 |
| **1.0 M NaHCO<sub>3</sub>** | 510 | 13.6 | 940 | 25 | 3190 | 85 |
| **2.0 M NaCl** | – | – | 890 | 47.2 | 720 | 38.4 |
| **0.15 M MgCl<sub>2</sub>·6H<sub>2</sub>O** | 40 | 0.15 | 30 | 0.12 | 80 | 0.33 |
| **0.5 M (NH<sub>4</sub>)<sub>2</sub>CO<sub>3</sub>** | 5 | 0.06 | 40 | 0.5 | – | – |
| **0.3 M CaCl<sub>2</sub>·2H<sub>2</sub>O<sup>1</sup>** | 5 | 1.5 | 0.5 | 0.15 | 2 | 0.6 |
| **1M HCl** | 80 | 1.1 | 100 | 15.6 | 50 | 8.4 |

<sup>1</sup>CaCl<sub>2</sub>·2H<sub>2</sub>O must be added fresh immediately before conducting each trial to avoid precipitation in storage.

Phase-specific electrolyte solutions were prepared by combining the master salt stock solutions with distilled water as shown in **Table 1** to reach target concentrations defined by INFOGEST 2.0 (12). These working electrolyte solutions were prepared in advance, aliquoted, and stored at −20°C. On the day of each experiment, aliquots were thawed before addition of CaCl₂, enzymes, or bile salts as required for each digestion phase.

### General Experimental Conditions

All phase-specific experiments were conducted independently in a final reaction volume of 1 mL and incubated at 37℃ to simulate physiological temperature. Reaction mixtures were prepared using the respective 1.25× simulated gastrointestinal electrolyte solution and phase-specific concentrations of CaCl_2_, digestive enzymes, and bile salts, as summarized in **Table 2**. The use of 1.25× electrolyte solutions allowed the simulated fluids to reach approximately 1× concentration following the addition of the other phase-specific components and bacteriophage suspension. Before phage exposure, 900 µL reaction mixtures were prepared, adjusted to the phase-specific pH, and pre-heated for 10 min. Enzyme stocks were prepared fresh on the day of each experiment and maintained at 4 °C or on ice during experimental setup to minimize enzymatic degradation. All amounts used were calculated to achieve the target conditions and final concentrations suggested by the INFOGEST 2.0 model (**Figure 1**).

**Table 2.** Composition of phase-specific reaction mixtures (recipe for 900 µL total volume).

|  | SSF Reaction | SGF Reaction | SIF Reaction |
| --- | --- | --- | --- |
| **Electrolyte Solution** | 800 μL<br>1.25× SSF | 799.5 μL<br>1.25× SGF | 800 μL<br>1.25× SIF |
| **0.3 M CaCl<sub>2</sub>** | 5 μL | 0.5 μL | 2 μL |
| **Enzymes** | 95 μL α-amylase<br>(790 U/mL) | 100 μL pepsin<br>(20,000 U/mL) | 48 μL pancreatin<br>(2100 U/mL) |
| **200 mM Bile Salts** | — | — | 50 μL |

Experiments were initiated by adding 100 µL of the working Sf14 suspension to the 900 µL phase-specific reaction mixture, yielding a final volume of 1 mL. Reaction tubes were briefly vortexed to ensure uniform mixing, and this moment was designated as the initial timepoint (t = 0). At each designated timepoint, 100 µL was transferred from the reaction tube into 900 µL of phage dilution buffer (PDB) to generate the first 10-fold serial dilution. Samples were immediately serially diluted to limit further exposure to experimental conditions. Viable phage particles were quantified by double-layer agar plaque assay and reported as PFU/mL.

Control reactions were prepared in parallel to each trial by combining 900 µL PDB with 100 µL of the same working Sf14 suspension and incubated under the same physiological temperature conditions. Control samples were collected at the final experimental timepoint to assess phage stability in the absence of simulated gastrointestinal conditions. Phages were serially diluted and quantified as described for the experimental groups. All experiments were conducted in at least 3 independent trials.

### Oral Phase

#### Salivary α-Amylase Stock Preparation

Although the INFOGEST 2.0 protocol indicates that salivary amylase is only necessary for the digestion of food containing starch, it was included in this study to more closely imitate oral conditions, as it is one of the main components that makes up 10-20% of saliva’s protein content (34). For the oral phase experiments, α-amylase (MP Biomedicals) was supplied as a lyophilized powder with a reported activity range of 500-1000 U/mg. For calculations, the midpoint of this range (750 U/mg) was assumed. The target enzyme activity in the final simulated salivary fluid (SSF) reaction mixture was 75 U/mL (12,13). To maintain a consistent final reaction volume of 1 mL, 95 μL of α-amylase stock solution was added to each reaction mixture. Delivering 75 units of activity within this volume required a stock activity of approximately 790 U/mL, which corresponds to a protein concentration of approximately 1.05 mg/mL based on the assumed enzyme activity. Accordingly, 2.1 mg of α-amylase powder was dissolved in 2 mL of 1× SSF electrolyte solution. The 1× SSF solution was prepared by diluting 1.25x SSF electrolyte with distilled water at a 4:1 ratio (1.6 mL 1.25× SSF combined with 0.4 mL distilled water). Although the INFOGEST 2.0 digestion method prepares α-amylase stock solutions in water, the enzyme was instead dissolved in SSF electrolyte adjusted to approximately pH 6.8 using small additions of 6 N HCl to achieve the enzyme’s optimal pH for its activity (13).

#### Oral Phase Infectivity Protocol

The SSF reaction mixture was prepared as outlined in **Table 2** and adjusted to pH 7 (range: 6.8 – 7.2). Immediately following equilibration of the SSF reaction mixture at 37℃, the oral phase was initiated upon phage exposure as described under *General Experimental Conditions*. Samples were collected at t = 0 min, 0.5 min, 1 min, and 2 min to represent the short residence time anticipated for the oral phase. Phages in the control PDB tube were sampled only at 2 min, corresponding to the final experimental timepoint.

### Gastric Phase

#### Isolated Effect of pH on Sf14 Infectivity

Since the gastric pH ranges from 1.5 – 3.5 while fasting (15), a preliminary test of PDB in the pH range of 2 – 3 was conducted first. Individual tubes containing 900 μL of PDB were adjusted to pH 2.0, 2.25, 2.5, 2.75, and 3.0 using small additions of 6 N HCl. Experimental and control tubes were set up as described above, with time points t = 0 min, 5 min, and 10 min up to pH 2.5, plus 30 min, 60 min, and 120 min for pH 2.75 and pH 3.0.

#### Pepsin Stock Preparation

Pepsin is the primary and only proteolytic enzyme present in the gastric phase, but its reported activity varies widely across studies due to differences in assay methods, unit definitions, and the presence of multiple enzyme isoforms (13). The INFOGEST 2.0 model recommends a final pepsin activity of 2,000 U/mL, indicating enzymatic activity rather than mass concentration. Pepsin (MP Biomedicals) was supplied as a powdered enzyme with an activity designation of 1:10,000. Since this format does not directly correspond to standardized enzyme units, the manufacturer’s data sheet was referenced, which reported activity based on a 10-minute assay, corresponding to approximately 1,000 U/mg.

To achieve the desired final activity, a pepsin stock solution of 20,000 U/mL was prepared by dissolving the appropriate mass of enzyme powder (20 mg per 1 mL of stock) in 1× simulated gastric fluid (SGF) electrolyte solution adjusted to an acidic pH (∼2.5). This allowed for the addition of 100 μL of stock to a 1.0 mL reaction mixture to achieve a final activity of 2,000 U/mL. Additionally, porcine pepsin is commonly used due to its high structural homology to human pepsin and its accessibility, making it a suitable model enzyme for simulated gastric conditions (13). The stock solution was prepared fresh on the day of each experiment and maintained on ice to preserve enzymatic activity.

#### Gastric Phase Infectivity Protocol

For the gastric phase, the electrolyte volume was adjusted slightly from 800 μL to accommodate the addition of CaCl₂ and pepsin while maintaining the consistent final reaction volume of 1 mL and preserving intended final concentrations of each component (**Table 2**, **Figure 4**). The pH of the SGF mixture was adjusted to 2–3 using small additions of 6 N HCl as needed. Experimental and control tubes were set up as described above, with time points t = 0 min, 5 min, 10 min, 30 min, 60 min, and 120 min.

### Intestinal Phase

#### Pancreatin Stock Preparation

Pancreatin (MP Biomedicals, CAS 8049-47-6), which is a mixture of pancreatic digestive enzymes, was used to simulate pancreatic enzyme activity in the intestinal phase. According to the manufacturer, the preparation contains a minimum protease activity of approximately 25 U/mg, along with additional amylase and lipase activities. The INFOGEST intestinal digestion model targets trypsin activity of approximately 100 U/mL in the final reaction mixture. Since the manufacturer specified total protease rather than trypsin activity, the reported protease activity was used to approximate the required pancreatin concentration. Each intestinal reaction mixture contained 48 μL of pancreatin stock within a total volume of 1.0 mL. To deliver approximately 100 units of protease activity in that volume, the stock solution needed to contain around 2,083 U/mL of activity. Based on the manufacturer-reported activity of 25 U/mg, this corresponds to a mass concentration of approximately 83.3 mg/mL pancreatin. For practical preparation, the concentration was rounded slightly upward to 84 mg/mL.

The stock was prepared by dissolving 84 mg of pancreatin powder in 1 mL of simulated intestinal fluid (SIF) electrolyte solution. The 1× SIF solution was obtained by diluting 1.25× SIF electrolyte with distilled water at a ratio of 4:1 (0.8 mL 1.25× SIF combined with 0.2 mL distilled water). When 48 μL of this stock was added to the intestinal phase reaction mixture, the resulting protease activity was approximately 100 U/mL in the final reaction volume.

#### Bile Salt Stock Preparation

Bile salts (Sigma-Aldrich, B8756) suitable for microbiological applications were used to simulate intestinal bile exposure with a final concentration of 10 mM. According to the manufacturer’s specification, this preparation contains approximately 50% sodium cholate and 50% sodium deoxycholate. Based on the molecular weights of these bile salts (430.6 g/mol and 414.6 g/mol, respectively), an approximate average molecular weight of 423 g/mol was used for calculations. To prepare a 200 mM bile salt stock solution, bile salts were dissolved in sterile, distilled water and vortexed until fully dissolved, then stored at 4°C. For each experimental reaction, this stock solution was vortexed and added to simulated intestinal fluid at a final bile salt concentration of 10 mM.

### Intestinal Phase Infectivity Protocol

Experimental and control tubes were set up as described under *General Experimental Conditions* and **Table 2**. The pH of the final intestinal mixture was measured using a calibrated pH meter and adjusted to approximately pH 7 (range: 7.0 – 7.4). Samples were collected and plated for quantification at timepoints t = 0 min, 60 min, 120 min, and 180 min. Control samples were quantified after 180 min to assess phage stability in the absence of electrolytes, intestinal salts, pancreatin, and intestinal pH.

## Acknowledgements

This work was supported by the National Institutes of Health award R01AI170608 to SMD. The funder had no role in study design, data collection and analysis, decision to publish, or preparation of the manuscript.

